# Audiovisual stimulation using wearable shutter glasses robustly evokes 40 Hz neuronal activity but does not modulate associative memory

**DOI:** 10.64898/2026.07.09.737418

**Authors:** Laura Hainke, Viktor Neumaier, Eleonora Marcantoni, Danying Wang, Kiera Capstick, Manuel Spitschan, James Dowsett, Simon Hanslmayr

**Author notes:** These authors contributed equally (Laura Hainke, Viktor Neumaier).

## Abstract

Audiovisual stimulation is a promising approach for studying and modulating human neuronal gamma (>30 Hz) oscillations and associated memory processes. Portable setups can increase ecological validity and therapeutic potential, but options remain limited. See-through shutter glasses are a new mobile technology that adds a visual flicker effect to what the user naturally sees. Here, we validated this method in a multisensory, cognitively relevant setting. We leveraged a previous experimental design from our lab, aiming to A) characterise the neuronal gamma activity evoked by shutter glasses, and B) conceptually replicate the previously reported effect of audiovisual gamma stimulation on memory accuracy, in line with a Spike Timing Dependent Plasticity model.

We recorded high-density Electroencephalography (EEG) from 24 healthy participants during an associative memory task. Video-sound pairs were presented with the sound amplitude-modulated at 40 Hz and 40 Hz visual flicker elicited by the shutter glasses, with a phase offset between both modalities. The visual flicker preceded the auditory modulation by 90 or 270 degrees. Participants were asked to remember the video-sound associations. They also underwent a visual-only condition and an electrically equivalent control condition. EEG evoked power and phase coherence were reconstructed at source level and analysed along with behavioural accuracy.

As expected, the shutter glasses robustly increased EEG evoked power and phase coherence at 40 Hz compared to the control condition. Effects were widespread and stronger than in a previous study not using shutter glasses. However, we did not replicate the previously reported effects of audiovisual phase offsets on memory accuracy. This could be due to reduced statistical power or methodological differences. Nonetheless, the validation of shutter glasses in a multisensory setting and the EEG analysis software, now improved and open source, enable important further investigations of audiovisual gamma stimulation in research and clinical settings.

## Introduction

Neuronal oscillations in the gamma band (>30 Hz) have been linked to memory processes in the human brain (1). Stimulating the brain at gamma frequencies can help investigate the mechanistic role of these oscillations in humans and modulate associated functions. Rhythmic sensory stimulation is a well-established, non-invasive method to do so: by presenting trains of stimuli – most commonly, visual and/or auditory – at a gamma frequency, neuronal activity can be induced or entrained at that same rate, and any effects on related memory functions can be assessed (2). One mechanism through which gamma oscillations may modulate memory is called Spike-Timing Dependent Plasticity (STDP): synaptic connections between neurons are thought to strengthen or weaken depending on the precise millisecond timing between pre- and postsynaptic spikes. Audiovisual gamma stimulation has yielded evidence for this mechanism in animals and humans (3,4). Moreover, the clinical potential of audiovisual gamma stimulation is increasingly being investigated for disorders marked by cognitive deficits, including Mild Cognitive Impairment and Alzheimer’s Disease (5–7). Beyond possible plasticity effects, gamma stimulation may - according to the glymphatic theory (8) - promote the brain’s fluid waste removal system and help clear neurotoxic molecules, such as parenchymal amyloid (9,10). Notably, audiovisual gamma stimulation has been proposed to be more effective than either modality alone (5,11).

Considering this broad potential, it is desirable to design audiovisual gamma stimulation protocols for use outside of standard laboratory environments, increasing ecological validity and therapeutic usability. While gamma-modulated auditory stimulation can be easily delivered via portable earphones, e.g., by playing modulated music, delivering visual stimulation at gamma frequencies is less straightforward. To date, clinical studies have mostly employed simple flickering lights, which patients look at for one hour per day (5). While simple to implement, this form of stimulation lacks meaningful visual content to keep users engaged. Moreover, cognitively relevant stimuli seem more efficient at propagating evoked gamma activity beyond the sensory brain areas (12,13), which seems desirable given that memory-relevant regions such as the hippocampus are located upstream from brain regions processing basic sensory input (14).

Shutter glasses address this challenge innovatively by modulating the visual field at any frequency of choice (15). They are liquid-crystal display glasses connected to a microcontroller that applies a low voltage to the crystal layer, switching the glasses between transparent and dark modes. This adds a full-field flicker effect to any real-world visual input the user currently perceives. Being portable and lightweight, they allow for stimulation during everyday activities. Recently, shutter glasses have been shown to elicit gamma responses effectively, demonstrating the necessary millisecond precision (16). However, this evidence is limited to low-density Electroencephalography (EEG) and a unimodal, visual-only stimulation protocol. This prompted us to investigate whether shutter glasses can efficiently modulate gamma activity in a multisensory setting with possible behavioural effects, and to explore how neuronal responses may differ from those elicited by standard computer-based visual flicker.

For the experimental design of this proof-of-concept study, we leaned on previous work from our lab conducted by Wang et al. (3), where audiovisual gamma stimulation yielded behavioural evidence for STDP in humans. Therein, participants were exposed to trials of concurrent soundtracks and videoclips modulated at 37.5 Hz. They were asked to remember the association between sound and video. During encoding, a phase offset was introduced: either the fluttering sound or the flickering video led by a few milliseconds. During recall, one group of participants was visually cued to re-watch a video and select the matching sound. Wang et al. showed that visual recall accuracy was higher when the 37.5 Hz-modulated video had led the modulated sound by 90° (6.6 ms) during encoding, potentially indicating strengthened connections from visual to auditory neurons, and lower when the sound had led by 90° (6.6 ms). The opposite pattern was reported for an auditory-cued group, in line with an STDP computational model. The visual cue group formed the basis for the present study.

The key aspect differentiating the present study from the original study by Wang et al. is the use of shutter glasses to modulate the visual input. Moreover, we opted for a modulation frequency of 40 Hz instead of the original 37.5 Hz, exploring whether the behavioural effect could be conceptually replicated at a frequency which is similar, but of higher clinical interest (5). To add further value beyond prior work, we also refactored the original MATLAB code for EEG source reconstruction and trial-level analysis by Wang et al., improving readability, reproducibility, and generalizability across multisensory stimulation experiments. The open-access pipeline is now freely available on GitHub (see Data availability).

The main goal of the present study was thus to characterise the neuronal responses evoked by shutter glasses at source level, compared to an electrically equivalent control condition and ‘standard’ monitor-based modulation used in the original study by Wang et al. We primarily expected that evoked power and Inter-Trial Phase Coherence (ITPC) would be higher in a unimodal visual flicker condition than in the control condition. Moreover, this study was designed as a conceptual replication of Wang et al. with a new technology for visual stimulation. Therefore, we compared EEG activity evoked by shutter glasses with the activity evoked by a standard computer monitor in the original study. Behaviourally, we hypothesized that recall accuracy would be higher in the realigned 90° (6.25 ms) phase condition compared to 270° (18.75 ms). In short, while we did not replicate the behavioural phase effect, the shutter glasses strongly and selectively modulated 40 Hz activity at source level, validating their use in further multisensory gamma experiments.

## Methods

### Sample

This study was approved by the local ethics committee of the School of Psychology and Neuroscience at the University of Glasgow. Volunteers were recruited via a university mailing list from 05.11.24 to 05.12.24 and compensated with 12 pounds per hour or course credit. We recruited 24 participants, matching the number of participants in Wang et al. who underwent EEG and were visually cued. We chose the visual cue condition over the auditory due to the stronger reported phase effect in the visual cue group (3). To further increase statistical power, we reduced the audiovisual phase offsets from four (0° / 90° / 180° / 270°) to the two phases relevant for the main effect (90° / 270°) while keeping the total number of trials equal. The participants were healthy, young adults (mean 23.17 years; range 18-31 years) with no history of or current neurological conditions, no current psychiatric disorders, normal or corrected vision, and normal hearing. Participants’ sex (13 female, 11 male) matched their reported gender; 3 were left-handed, and 21 were right-handed.

### Protocol

After obtaining participants’ written informed consent, the EEG system was set up. Participants sat 60 cm away from the monitor, with their head on a chinrest, in an electrically shielded and illuminated booth. They wore EEG-compatible insert earphones and the shutter glasses (Fig. 1A) for visual stimulation.

**Fig. 1:**
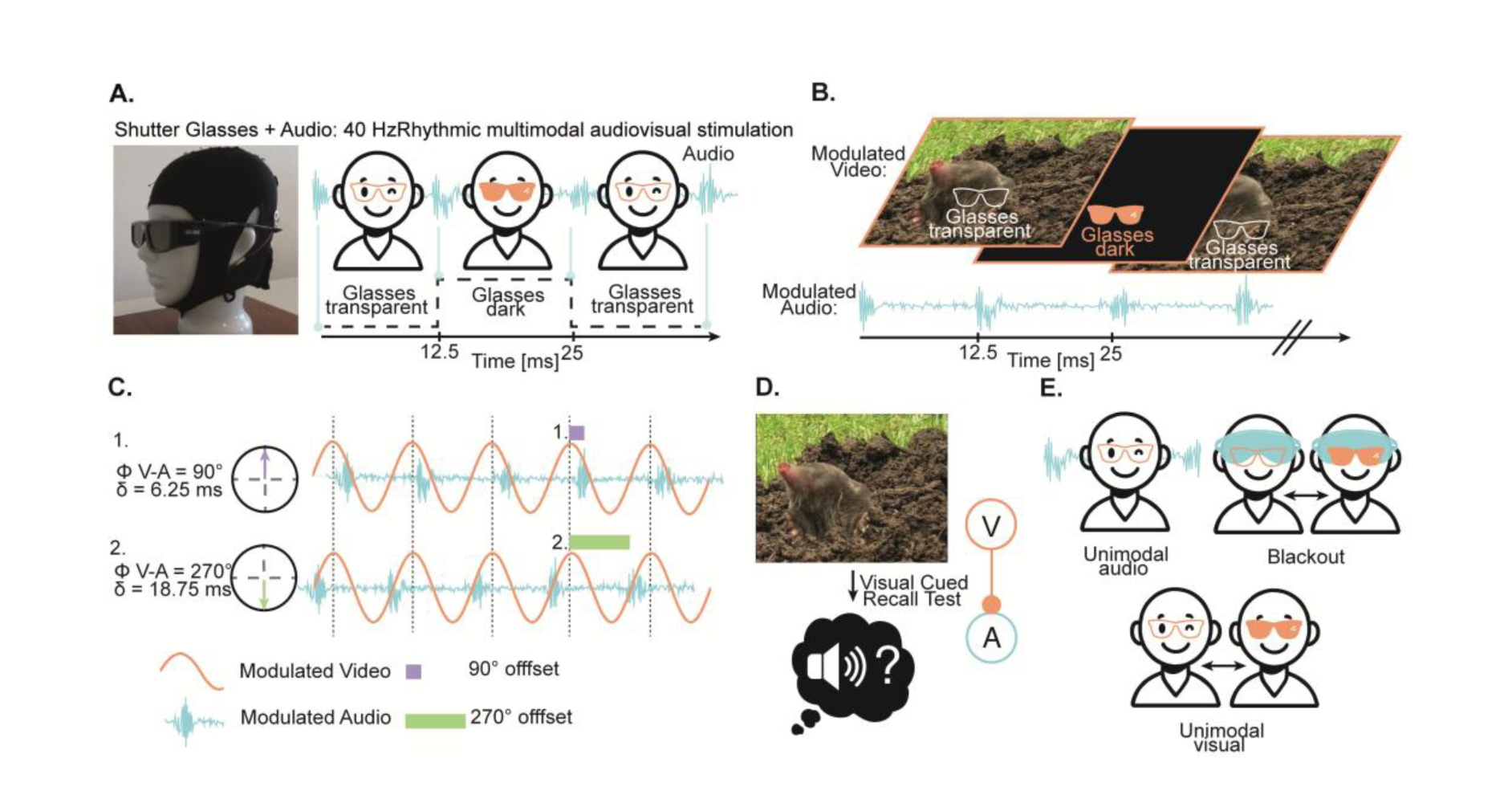
Experimental protocol. (A) left: shutter glasses with exemplary EEG cap on head dummy, right: Timing demonstration of shutter glasses changing from transparent to dark. (B) Exemplary video frames that were viewed during encoding phase wearing the shutter glasses, with a concurrent audio clip. (C) The amplitude of sounds and the luminescence of the video via shutter glasses were both modulated at 40 Hz with two phase offsets 90 and 270. The green and purple bars represent peak delays between a video (flicker leading) and a sound (flutter trailing). (D) In the memory test phase, the video was presented again; participants were asked to select the corresponding sound. (E) The three unimodal conditions: audio-only, visual-only, or blackout. The latter was a control condition in which participants wore a cloth mask underneath the flickering shutter glasses.

Following the design by Wang et al., the main part of the experiment started with four practice trials. 16 blocks of an associative memory task followed, each with 12 trials. Each block included an encoding, a distractor, and a recall phase. During encoding, 12 randomised sound-video pairs were presented for 3 s each. Unlike in Wang et al., the stimulus modulation frequency was 40 Hz, and the flicker was elicited by the shutter glasses, not the PC screen (Fig. 1B). Participants were instructed to remember the association between video and sound, and to rate how well they match from 1 (the sound does not match the contents of the movie at all) to 5 (the sound suits the movie very well) using keyboard numbers. Of the 12 trials, half were randomly assigned to the 90° condition and half to the 270° condition (i.e., the modulated audio track was delayed in relation to the video and flicker by 90 or 270 degrees, corresponding to 6.25 or 18.75 ms; Fig. 1C). A fixation cross was displayed between trials, with jittered inter-trial intervals (ITIs) always ranging from 1 to 3 s throughout the full experiment. In the distractor phase, a random number between 170 and 199 was shown, and participants were asked to count backwards in steps of 3 for 30 s. Finally, during recall, the 12 video clips presented in the encoding phase were shown in a random order. For each of those, participants were prompted to choose the matching sound out of 4 options (Fig. 1D).

In the next part (Fig. 1E), participants underwent 50 trials of a unimodal auditory condition. As in Wang et al., soundtracks were presented for 3 s with jittered ITIs, and subjects rated their pleasantness on a scale from 1 (very unpleasant) to 5 (very pleasant). The participants wore the glasses, which remained transparent, while looking at a fixation cross. 50 trials of a unimodal visual condition followed, in which videos and flicker were presented for 3 s with jittered ITIs. Participants were asked to rate their pleasantness while wearing the earphones without sound.

Lastly, participants underwent a “blackout” control recording. Here, they wore a black cloth mask underneath the shutter glasses, preventing sight, as well as the earphones without sound. 50 trials were recorded, in which the same randomised videos as in the unimodal visual condition were displayed consecutively, and the shutter glasses flickered concurrently, for 3 s per trial with jittered ITIs. While the participant was unable to perceive the videos or flicker, this ensures that this recording, serving as a baseline and control, is electrically equivalent to the unimodal visual condition. After 2.5–3 h, the experiment was terminated.

### Materials

EEG was recorded using a 128-channel BioSemi ActiveTwo system (BioSemi B.V., Amsterdam, Netherlands). Data were sampled at 4096 Hz using the BioSemi ActiView software. Electrode positions for each participant were tracked using a Polhemus Space3 Fastrak device (Colchester) and recorded using Brainstorm3 (17), implemented in MATLAB (R2022b; MathWorks, Natick, MA, USA).

Stimuli were presented using the Psychophysics Toolbox extensions (18–20) in MATLAB (R2022a), as in Wang et al. The videos were displayed on a high-performance VIEWPixx /EEG monitor (VPixx Technologies Inc., Quebec, Canada) with a 120 Hz refresh rate. The audio was presented via insert earphones (ER-3C, Etymotic Research). Millisecond accuracy of stimulus presentation was verified before study start using the Black Box ToolKit v3™ (BBTK v3; The Black Box ToolKit Ltd., England, UK). The same neutral video clips and soundtracks from different categories were used as in Wang et al., but the video clips had a frames-per-second rate of 40 and were not modulated.

Instead, to elicit a visual flicker effect, we used custom-built shutter glasses (15). They are adapted liquid-crystal display glasses (SainSonic, Texas, US), battery-powered and connected to a microcontroller (Arduino Uno, Scarmagno, Italy). When a low voltage is applied across the liquid crystal layer within the glass, it becomes darker; if this voltage is uniformly applied to both glasses at a chosen frequency, alternating between transparent and semi-opaque with switch times <1 ms, this elicits a visual flicker effect. Of note, this modulates the full visual field, not only the computer screen.

We used 40 Hz as the modulation frequency for visual and auditory input instead of the original 37.5 Hz for two main reasons: first, a proof-of-concept for the modulation of gamma activity using shutter glasses is most useful at 40 Hz, since this is the frequency most widely employed in the (sub)clinical literature (5). Second, with 40 being a multiple of the monitor’s 120 Hz refresh rate, this prevents interference between the two visual sources at an unintended frequency. We also used 40 Hz as the stimulation frequency in the unimodal conditions, unlike the 4 Hz used in Wang et al.’s unimodal conditions. This allowed us to assess whether the shutter glasses effectively elicited gamma activity in a visual-stimulation-only condition, compared to an electrically equivalent control condition – which was central to the present proof-of-concept study – without increasing participant burden.

In the combined audiovisual tasks, both the soundtracks’ amplitude and the glasses’ transparency were modulated at 40 Hz. The visual flicker, starting in synchrony with the videos in a transparent state, had a square-wave pattern and a duty cycle of 50 % (12.5 ms transparent, 12.5 ms dark). The soundtracks, modulated with a sine wave and starting at maximum volume, were presented concurrently with the videos, with a delay of 40 ms to compensate for faster neuronal auditory processing compared to visual processing (21). Additionally, the fluttering auditory track was phase-delayed by either 90° (6.25 ms) or 270° (18.75 ms) relative to the visual flicker. The sound and video contents were unrelated; pairings were counterbalanced across phases and participants.

### EEG preprocessing

EEG data were preprocessed using the Fieldtrip toolbox (22), akin to Wang et al. (3). First, data were epoched from 2 s pre-stimulus to 5 s post-stimulus onset. Stimulus onset times were defined based on triggers from the shutter glasses. Data were bandpass-filtered between 1 and 120 Hz and bandstop-filtered between 48-52 and 98- 102 Hz to remove line noise, then downsampled to 512 Hz. Bad channels and trials with large artefacts were removed by visual inspection before Independent Component Analysis was applied to remove ocular and cardiac artefacts. Bad channels were then interpolated using nearest-neighbour triangulation based on individual electrode positions. Lastly, after average re-referencing, any trials with remaining artefacts were manually rejected. The remaining clean trials were subjected to further analysis.

### Source reconstruction

Following the pipeline by Wang et al., we first estimated the origin of neuronal responses to 40-Hz-modulated audiovisual stimuli using data from the unimodal visual and auditory conditions. This facilitates the localisation of otherwise correlated neuronal sources by Linearly Constrained Minimum Variance (LCMV) beamforming. We then reconstructed the data acquired during multimodal stimulation at the localised auditory and visual sources (3) (Fig. S1-3).

First, individual source models were prepared using individuals’ digitised electrode positions aligned to a template head model, along with a template volume conduction model (23). The procedure then differed slightly between the unimodal auditory and visual conditions: For the unimodal visual condition, electrical potential time series from 2019 virtual electrodes – uniformly distributed within a participant’s modelled 3D head space – were estimated through LCMV beamforming, using leadfields based on scalp potentials. For the unimodal auditory condition, data were first transformed to Scalp Current Density (SCD) using the finite-difference method. The same was done to the leadfields by applying the matrix used for the sensors’ SCD transformation. This additional step prior to LCMV helps the beamformer localise two correlated, spatially separated auditory sources (24). Then, for both unimodal conditions alike, data were averaged across trials at each virtual electrode and subjected to time-frequency analysis.

As in Wang et al., the evoked power (see Outcomes) at each virtual electrode was averaged over 0.75-2.75 s post-stimulus onset, within ±0.5 Hz of the frequency of interest (here, 39.5 Hz – 40.5 Hz). Evoked power must be normalised to be compared across participants; however, the method previously used by Wang et al. – randomly shifting each trial forward by 0, 90, 180, or 270° before averaging to create a shuffled baseline – proved suboptimal for this dataset, as it resulted in implausible auditory sources. Moreover, source localisation results were unstable, varying considerably within participants depending on the randomisation seed used. We therefore opted for the pre-stimulus period (-1.75 to -0.25 s before stimulus onset) of the averaged trials as the baseline here, due to its deterministic nature and common use in the literature (25). The average evoked power at each virtual electrode was thus normalised by subtracting the baseline evoked power and dividing by its average.

Finally, for each unimodal condition, the normalised evoked power at each virtual electrode was grand-averaged across all participants and interpolated to the MNI magnetic resonance image template (26). The final coordinates for the Regions of Interest (ROIs) to be used across subjects in further analyses – one visual source and two auditory sources, one per hemisphere – were the locations where the grand-average evoked power was estimated to be maximal across participants in the respective unimodal condition.

Having determined the visual and auditory ROIs in the present sample, we again followed the procedure of Wang et al. to estimate time-series data at the ROI sources during multimodal stimulation using LCMV beamforming. For the reconstruction at the two auditory ROIs, first, the sensor-level data from the multimodal condition were SCD-transformed; then, one set of spatial filters per hemisphere was computed with LCMV; finally, the two sets of spatial filters were applied to the SCD-transformed time series. Time-series data were thereby estimated at the left and right auditory ROIs (24). The time-series data at the visual ROI were reconstructed directly with LCMV without prior SCD transformation, akin to the unimodal procedure. As beamforming results are inherently ambiguous regarding dipole orientations, the signs of the reconstructed time series were manually adjusted (see (3)). Lastly, the reconstructed time series data from both auditory ROIs were averaged.

### Trial label realignment

As Wang et al. showed (3), the phase difference between time series at the visual and auditory ROIs often does not equal the experimentally defined phase offset between visual and auditory stimulus trains, likely due to individual differences in transduction delays. Trials must therefore be relabelled based on the actual neuronal phase difference between the visual and auditory ROI time series, bandpass-filtered between ±2.5 Hz around the stimulation frequency for better detection of gamma signals (here, 37.5 - 42.5 Hz).

At the single-trial level, Wang et al. did so by calculating the instantaneous phase differences between the visual and auditory ROIs, computing the mean angle across data points 0.5 – 2.5 s post-stimulus-onset, sorting the trials by mean angle values from -pi to pi, and then dividing the trials into four equally sized bins to relabel them (3). However, in our dataset, the actual phase offsets were less evenly distributed. For more accurate relabelling, we restricted the four phase bins through angle boundaries of ±45° around the phase of interest: e.g., for a trial to be assigned to the 90° phase bin, its mean phase difference between visual and auditory ROIs’ time series had to be >45° and <135°. Trials falling into 0° or 180° bins were rejected in our analyses.

### Outcomes

At the electrophysiological level, the two main outcomes were evoked power and Inter-Trial Phase Coherence (ITPC) at the stimulation frequency of 40 Hz. For analyses, time-frequency representations were obtained in the range of 39–41 Hz and -2–4 s. We used single-taper convolution based on Slepian sequences, a sliding window of 1 s, and a spectral smoothing bandwidth of 2 Hz. Evoked power values were baseline-normalised as described above and then averaged over frequencies (39.5–40.5 Hz) and timepoints (0.75–2.75 s post-stimulus) of interest. To compute the ITPC, the complex Fourier spectrum of each trial was normalised by its magnitude. The resultant vector length was computed across trials for each time point. The values averaged across frequencies (39.5–40.5 Hz), and timepoints (0.75–2.75 s post-stimulus) of interest constitutes the ITPC. At the behavioural level, the main outcome was recall accuracy (correct or incorrect) for each video-sound association.

### Statistical analyses

For paired comparisons, we used one-sided paired t-tests or Wilcoxon signed rank tests if parametric assumptions were violated. For unpaired comparisons, we used t-tests or Mann-Whitney U tests, as appropriate. A one-sample t-test was used to compare individual peak voxels in the present sample to the original study’s group peak voxel. For recall accuracy as a binary variable, a Generalised Linear Mixed Model (GLMM) for binomial distributions was computed using a Nelder Mead optimizer. Lastly, to assess any relationships between electrophysiological variables and recall accuracy, Pearson correlations were computed. The significance threshold was α=0.05 and p-values were False-Discovery-Rate (FDR)-corrected where appropriate. Participants with <5 trials in any tested condition were excluded from the respective test.

## Results

First, we inspected the origins of neuronal responses to audiovisual stimulation estimated by the source reconstruction pipeline. In the unimodal auditory condition, 40 Hz activity was most prominently evoked bilaterally in posterior temporal regions (Fig. 2). The unimodal visual condition revealed an occipito-parietal source of evoked 40 Hz activity. Compared to Wang et al., the detected source in this study appeared more dorsal (Fig. 2). This was confirmed by means of a one-sample t-test comparing the z-coordinate of each participant’s peak visual source to the z-coordinate of the group peak voxel from (3) (*N*=24; *p*=.006; *t*=2.74; *r*=0.59; 95% CI [0.27;1.00]).

**Fig. 2:**
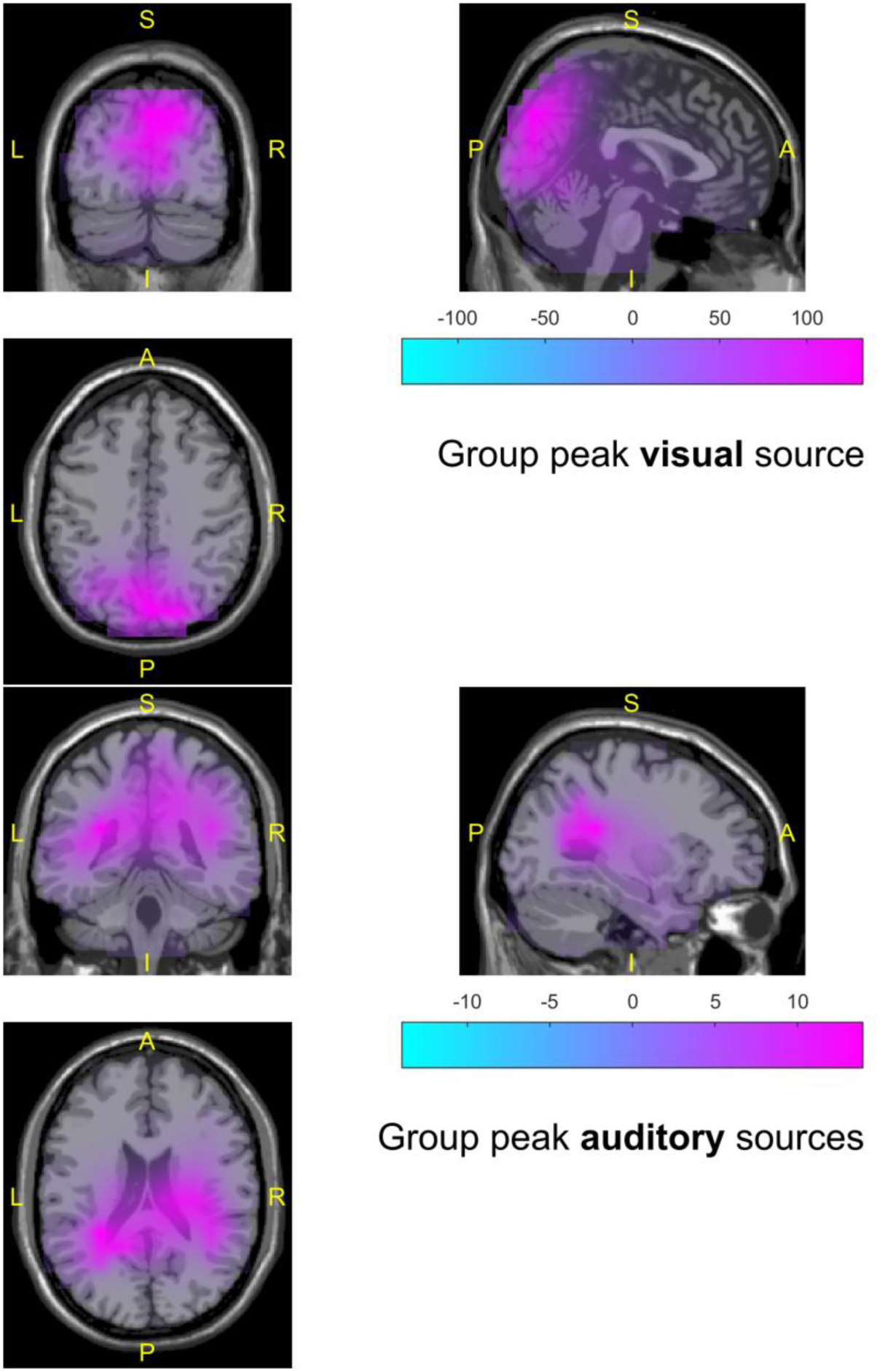
40 Hz sensory activity was predominantly evoked at occipito-parietal and posterior temporal regions. Baseline-normalised evoked power grand-averaged across participants and interpolated to the MNI template reveals the group peak visual and auditory sources.

Confirming our primary hypothesis, the shutter glasses clearly and robustly modulated 40 Hz EEG activity. At the reconstructed visual source, both normalised evoked power and ITPC at 40 Hz were significantly higher in the unimodal visual condition than in the electrically equivalent blackout control condition. Wilcoxon signed-rank tests revealed strong effects for evoked power (blackout median=0.632, SD=1.099; unimodal visual median=73.366, SD=128.701; *V*=276, *p*<.001, *r*=1, 95% CI [1;1]; Fig. 3A) and ITPC (blackout median=0.156, SD=0.29; unimodal visual median=3.954; SD=2.35; *V*=276, *p*<.001, *r*=1, 95% CI [1;1]; Fig. 3B).

**Fig. 3:**
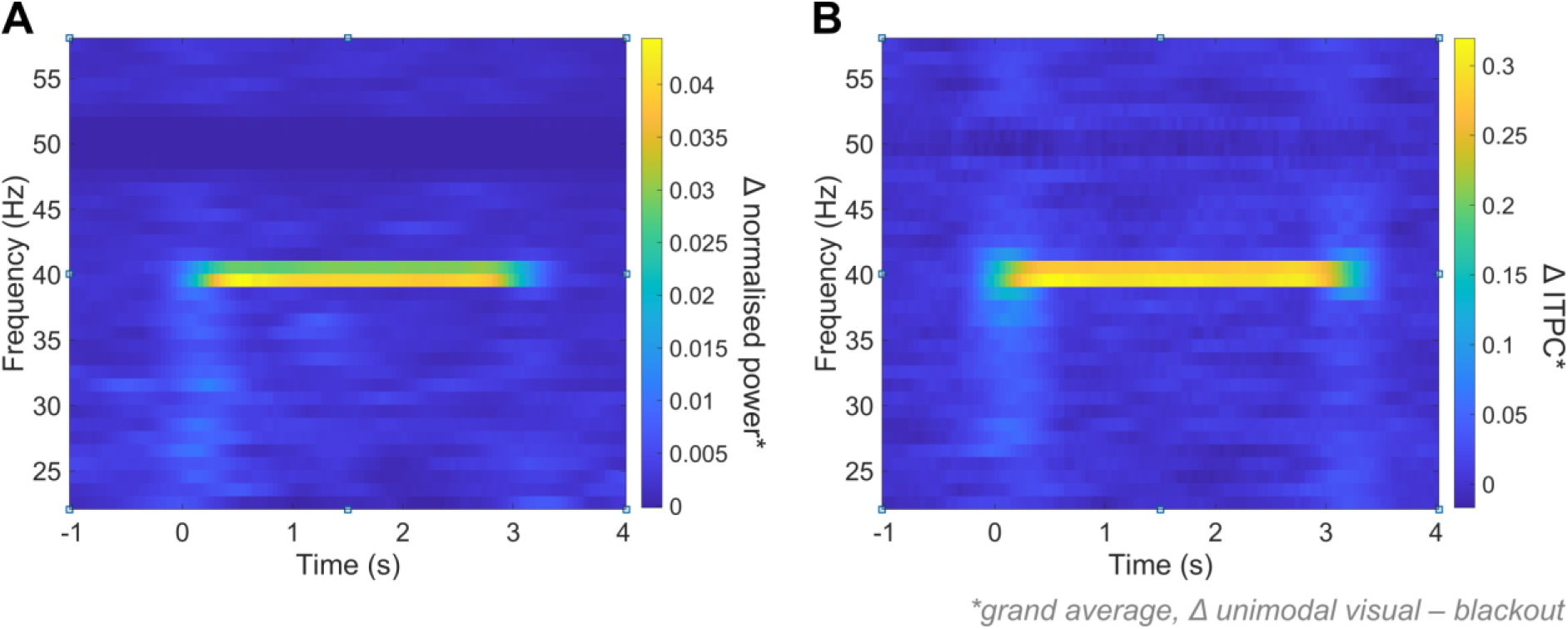
Grand-average evoked power and ITPC at 40 Hz were significantly higher in the unimodal visual than in the control condition. At the reconstructed peak visual source, (A) baseline-normalised evoked power and (B) Inter-Trial Phase Coherence at 40 Hz were significantly higher in the unimodal visual condition than in the electrically equivalent blackout control condition; only at the frequency and times of interest (frequency range expanded here for visualization purposes). Times relative to audiovisual stimulation onset.

This finding is supported by a significant response across all 128 electrodes – compared to blackout, for both power and ITPC, and only during the stimulation period (for all electrodes, FDR-corrected *p<*.05). This pattern demonstrates strong, widespread activation selective to the stimulation period (Fig. 4).

**Fig. 4:**
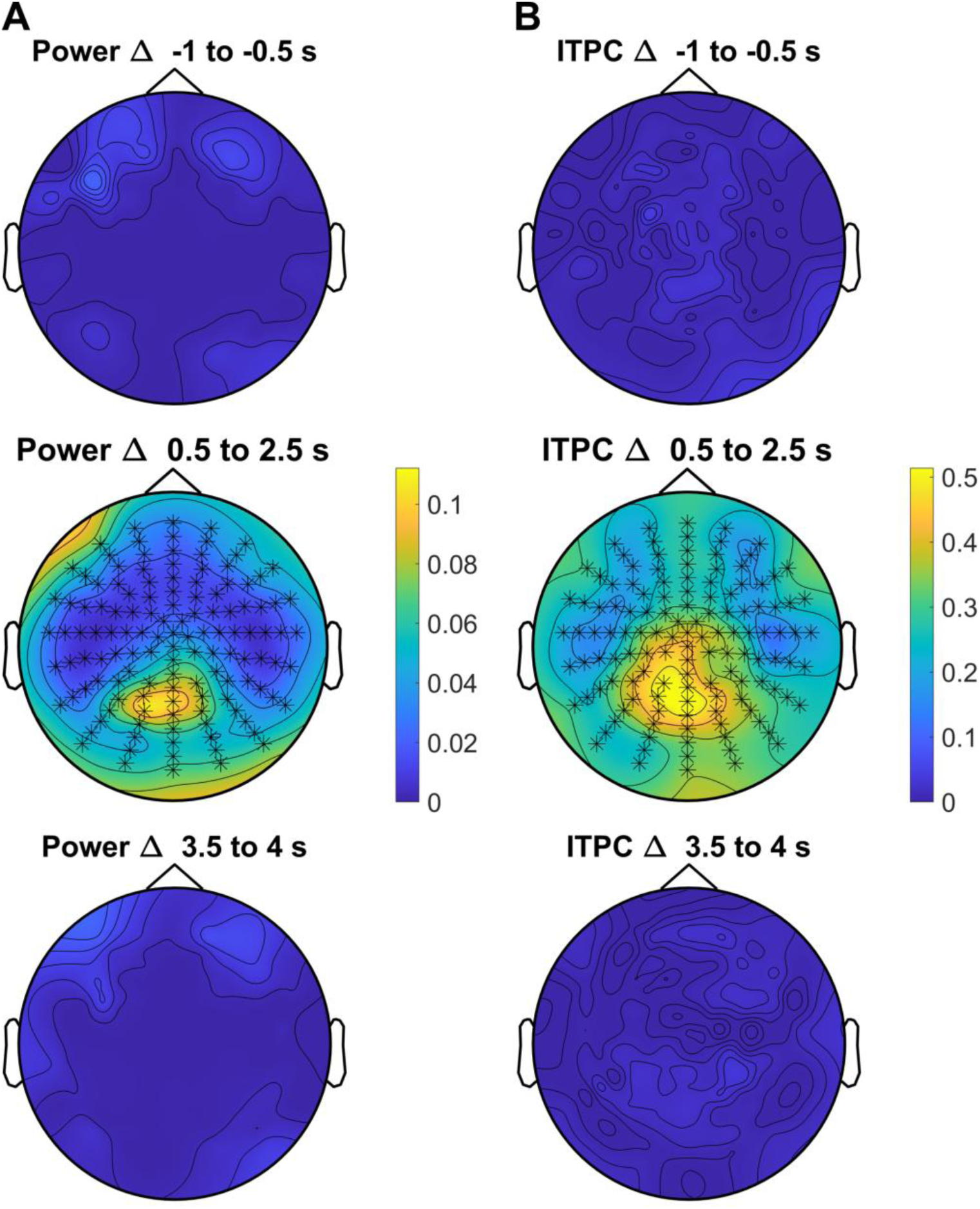
Shutter glasses yield widespread, coherent 40 Hz activity. During visual stimulation, but not in pre- and post-trial periods, normalised evoked power (A) and Inter-Trial Phase Coherence (B) at 40 Hz were significantly increased across all electrodes after False-Discovery-Rate correction. Values represent the difference between the unimodal visual and the control condition, ruling out electrical artefacts.

Next, we tested whether the evoked power and ITPC at 40 Hz at our reconstructed peak visual source differs from the values at 37.5 Hz in Wang et al.’s data, at their respective visual source. The same baseline normalisation method was applied to both datasets. The comparison was performed for multimodal data, where the difference in stimulation frequencies between studies was smallest, using Mann-Whitney U tests. Indeed, we observed a stronger activation in our sample, both in evoked power (Wang et al. median=92.82, SD=327.43; here, median=422.39, SD=779.17; *W*=131, *p*=.001, *r*=0.47, 95% CI [0.2;0.7]; Fig. 5A) and ITPC (Wang et al. median=0.44, SD=0.17; Hainke et al. median=0.76, SD=0.23; *W*=107, *p*<.001, *r*=0.539, 95% CI [0.28;0.73]; Fig. 5B). Note that the total number of trials per participant was significantly higher in the present study (Wang et al. mean=140, SD=26; here, mean=172, SD=22; *t*=-4.70, *p*<.001).

**Fig. 5:**
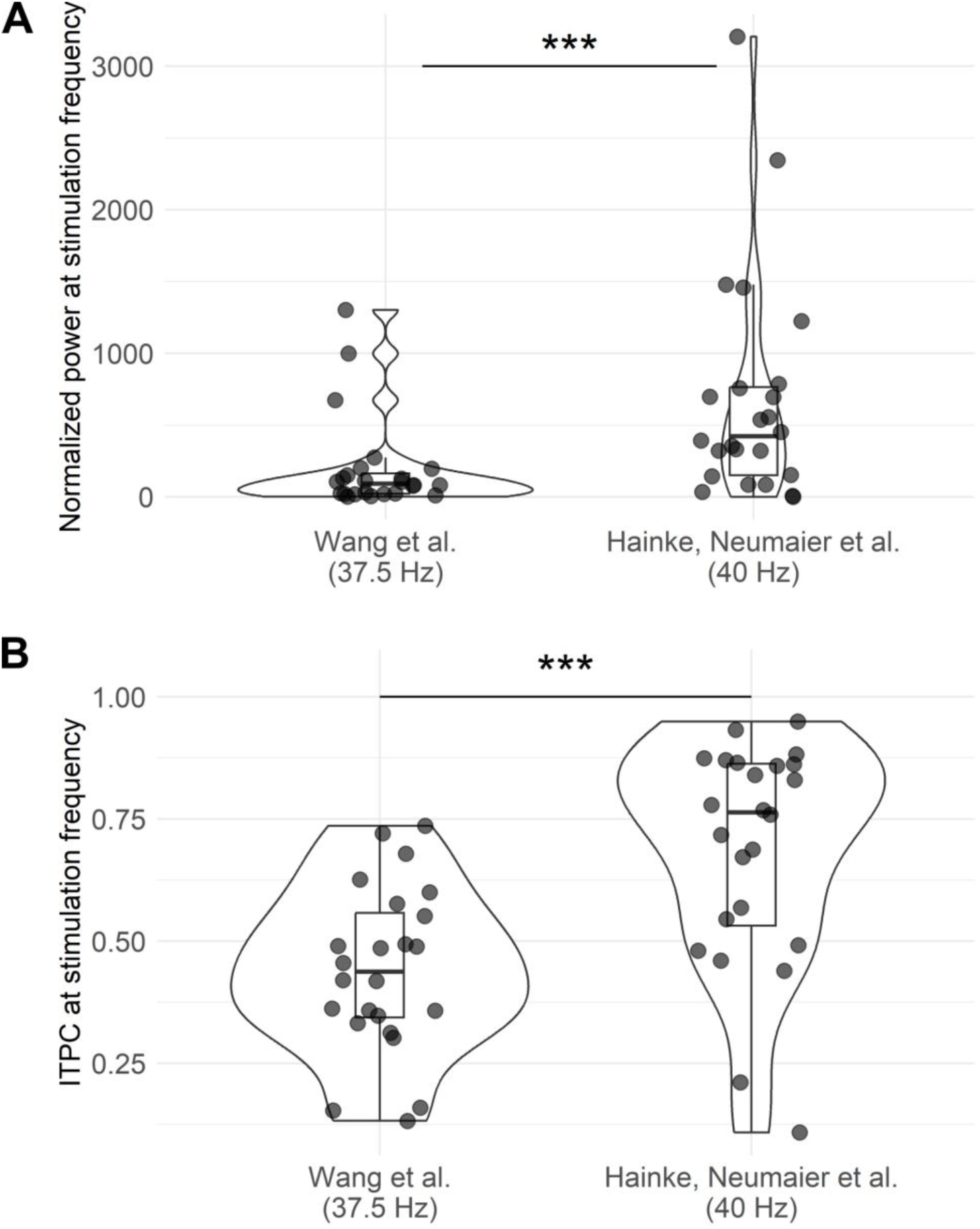
Shutter glasses evoke stronger and more coherent gamma activity than a flickering computer screen. (A) Comparison between Wang et al.’s and the present study, regarding normalised evoked power. (B) Comparison between Wang et al.’s and the present study, regarding Inter Trial Phase Coherence (ITPC). *** for p<.001. Boxplots show medians and interquartile ranges.

As for the main behavioural hypothesis, we first confirmed that trial realignment as in Wang et al. was successful. Each trial of each participant was reassigned to one of four phase bins (0/90/180/270°) based on the mean neuronal phase offset between the reconstructed visual and auditory sources (see *Trial label realignment*). For the trials assigned to the 90° and 270° phase bins, the reconstructed and bandpass-filtered time series were averaged within and across participants for the visual and auditory sources, respectively, yielding the grand average event related potentials shown in Fig. 6A-B. The mean phase offsets of these visual and auditory grand average time series closely reflect the intended offsets of 90° and 270°, respectively, confirming correct realignment at group level.

**Fig. 6:**
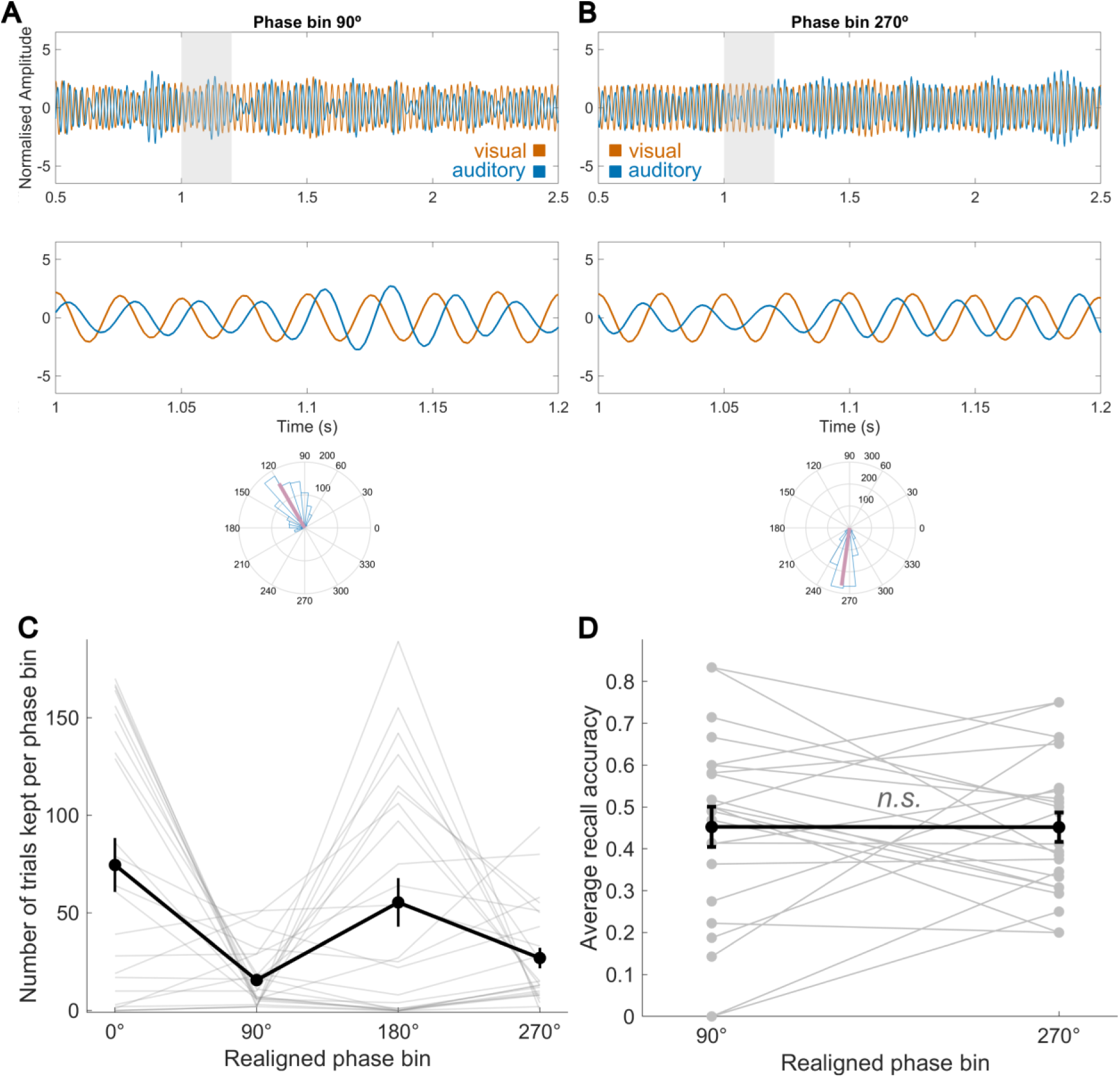
Trial realignment based on actual neuronal offsets. (A) Grand-average, bandpass-filtered event-related potentials for the reconstructed visual (orange) and auditory (blue) sources after realignment to the 90° phase bin, for the full trial period and zoomed in. The circular plot shows the audio-visual phase difference at source level in degrees, confirming an average value near 90°. (B) Equivalent to A, for the realigned 270° phase bin. The circular plot again confirms successful trial realignment, with an average phase near 270°. (C) The number of trials retained per participant and bin after trial realignment. (D) Average recall accuracy values within and across participants, for the two phase bins of interest, after trial realignment. In C and D, light grey lines represent participant-level data, black lines mark group means, and whiskers represent the standard error of the mean.

Despite successful realignment, we could not replicate the higher recall accuracy in the realigned 90° phase bin compared to 270°. A binomial GLMM with accuracy as the outcome, realigned phase as the fixed effect, random intercepts for phases and random slopes for participants showed no significant effects (all *p*>.05) and was not superior to an intercept-only model (*p*>.05). Note that although we presented an equal number of trials to each participant with an experimentally manipulated audiovisual offset of 90° or 270°, because trials were relabelled to 0/90/180/270° bins based on reconstructed neuronal phase offsets, the final number of trials per bin differed. Unfortunately, the final number of trials per bin was very uneven, and the phase bins of interest (90° and 270°) were left with a particularly low number of trials (Fig. 6C). This resulted in the exclusion of 12/24 participants with <5 trials in the 90° and/or 270° phase bin, substantially lowering the statistical power of this test (Fig. 6D) and of any subsequent tests related to phase-realigned trials.

Two further findings originally reported by Wang et al. could not be replicated: In our sample, the ITPC at the visual ROI was not significantly higher in the realigned 90° condition (paired t-test *p*>.05). The difference in ITPC at the visual ROI between 90° and 270° conditions did not significantly correlate with the difference in recall accuracy between 90° and 270° conditions either (Pearson correlation *p*>.05). However, given the low number of trials in these conditions after realignments, these results should not be interpreted further. Lastly, we explored if evoked 40 Hz power at the visual ROI during multimodal encoding might correlate with overall recall accuracy, but a binomial GLMM with accuracy as the outcome, evoked 40 Hz power at the visual ROI as the fixed effect, and participant-level slopes did not show significant results (*p*>.05).

## Discussion

In this study, we delivered 40 Hz audiovisual stimulation at different phase offsets using see-through shutter glasses, assessing effects on EEG activity and the recall accuracy of modulated video-sound pairs. We demonstrated that shutter glasses evoke strong, widespread gamma activity, paving the way for this technology’s use in a range of multisensory applications. Although we could not replicate the behavioural effects reported by Wang et al., we contributed to the field by probing the parameter space of this effect and by sharing an improved, reproducible analysis pipeline for high-density EEG source reconstruction of multisensory time series.

The primary aim of this study was to validate the use of shutter glasses in a multisensory setting. The glasses clearly evoked EEG activity specific to the stimulation frequency (40 Hz) and duration, significantly detectable at all 128 electrodes. Importantly, artefacts induced by the shutter glasses were ruled out as a possible confounder through the electrically equivalent “blackout” control condition. Interestingly, both evoked power and ITPC were higher than in Wang et al.’s study. The larger number of trials in our sample could partially account for the increased evoked power, possibly by increasing the signal-to-noise ratio. ITPC, on the other hand, tends to decrease with more trials (27). It is therefore plausible that the shutter glasses’ visual modulation and triggers were more precisely timed. Given that a 40 Hz gamma oscillation cycle is only 25 ms long, any millisecond-scale improvements in timing precision can influence outcomes. Moreover, the glasses’ full-field visual flicker effect likely drove both the magno- and parvocellular visual pathways (28), increasing the number of activated neurons. Lastly, the frequency of 40 Hz, possibly closer to participants’ resonant gamma frequencies (29), may also have contributed to a larger synchronised neuronal population, increasing coherence.

Regarding the behavioural outcomes, the advantage in memory performance at a 90° audiovisual offset reported by Wang et al. could not be replicated in this sample, for which there may be several reasons. One possibility is the lack of statistical power after trial realignment based on actual neuronal offsets. Wang et al. flipped the 90° (6.6 ms) and 270° (20 ms) condition labels for their analyses, as most experimental 90° trials resulted in an actual neuronal offset of 270°, and vice versa. Therefore, in the present experiment, only the two phases of interest (90°, 6.25 ms, and 270°, 18.75 ms) had been presented to participants instead of the four original phases while keeping the total number of trials constant, in an attempt to increase statistical power. However, in the present sample, most trials showed actual neuronal offsets of 0° and 180°, resulting in the exclusion of 12/24 participants in behavioural analyses and greatly limiting statistical power. This could be due to individual neuronal variability in visual and/or auditory processing delays (30), nonlinear audiovisual interactions (31), or gamma-dependent, trial-level temporal recalibration of asynchronous audiovisual stimuli (32). Future experiments should ideally present four or more phase offsets.

Moreover, individual brain source voxels showing the strongest visual activation (and, consequently, the group-level peak voxel) were significantly more dorsal in our sample compared to Wang et al. 2023. This may have resulted from a more widespread activation of the parvo- and magnocellular pathway by the shutter glasses’ full-field contrast and may also have contributed to the discrepancy in behavioural results. The dorsal visual pathway receives substantial input from the magnocellular pathway (28), which is why the source in the present study may be more dorsal compared to Wang et al., where the periphery was not flickering. The different stimulation frequencies in the unimodal conditions (4 Hz in (3) vs. 40 Hz here), which we chose here to validate the shutter glasses in both uni- and multimodal conditions alike, may also have played a role.

Since evoked visual power at this ROI did not correlate with recall accuracy, it may be less relevant for memory processes than the ROI identified in Wang et al.’s sample. It is also possible that the 37.5 Hz modulation frequency used by Wang et al. is more conducive to memory effects than the commonly used 40 Hz. Further research on individual gamma frequencies and how they relate to memory, particularly when coupled with theta oscillations, is needed (33,34). Overall, we argue that the behavioural null results reported here do not necessarily discredit the original findings (3). They do, however, suggest that the effect is sensitive to certain boundary conditions such as the number of trials, the exact gamma stimulation frequency, and the spatial location of the peak neuronal sources.

The EEG source reconstruction code used in this study, adapted from Wang et al. for better reproducibility, readability, and generalisability, is another important contribution of the present study to the field. Multisensory stimulation and its effects on memory are also being explored at other frequency bands (21,35,36), and for better study comparability, it is important to harmonise analyses and outcomes. The present open-access pipeline can support this process. In particular, reconstructing not only the most active neuronal sources but also their correlated time series poses a challenge for common source reconstruction algorithms like the beamformer. The present pipeline addresses this challenge by building on prior methodological advances (24). While naturally based on the previous work by Wang et al., the refactored code has been restructured and documented for reuse in diverse multisensory memory experiments, providing a useful resource for future research.

To conclude, the present study contributes to the fields of Cognitive Neuroscience and Neurotechnology by validating the use of see-through shutter glasses as a novel technology for gamma neuromodulation in multisensory settings. We demonstrated that this form of visual stimulation evokes widespread neuronal gamma activity, potentially yielding stronger evoked power and phase coherence than a standard computer screen, while enabling portable use outside the lab. Due to its high usability and ecological validity, this technique has the potential to boost progress in multisensory gamma neuromodulation for clinical purposes (5,7,37). Furthermore, we have identified relevant parameter boundaries of the memory effect reported by Wang et al. (3), countering publication bias. Lastly, the validated shutter glasses and the open-access EEG source-reconstruction pipeline enable a range of further experiments in this field, for example, on theta-gamma coupling, long-term at-home interventions, or populations with memory impairments.

## Acknowledgments

None

## Data and code availability

Upon publication, the preregistration and anonymised data will be made available on Open Science framework (osf.io). The source reconstruction pipeline, reworked for use across multimodal RSS studies, is now available on GitHub under the following link: https://github.com/NoT-CoOLab/audiovisual-RSS-pipeline The code and outputs pertaining to this specific study can also be found on GitHub under the following link: https://github.com/laura-hainke/Hainke-Neumaier-et-al-2026

## CRediT contributions

Conceptualization – LH, JD, SH

Data Curation – LH, VN

Formal Analysis – LH, VN

Funding Acquisition – LH, SH

Investigation – LH, VN, KC

Methodology – LH, VN, EM, JD, DW

Project Administration – LH, SH

Resources – JD, SH

Software – LH, VN, EM, DW

Supervision – JD, MS, SH

Validation – LH, VN, EM, KC

Visualization – LH, VN, DW

Writing – Original Draft Preparation – LH, SH

Writing – Review & Editing – LH, VN, EM, MS, SH, KC, DW, JD

## Supplementary Materials

**Fig. S1:**
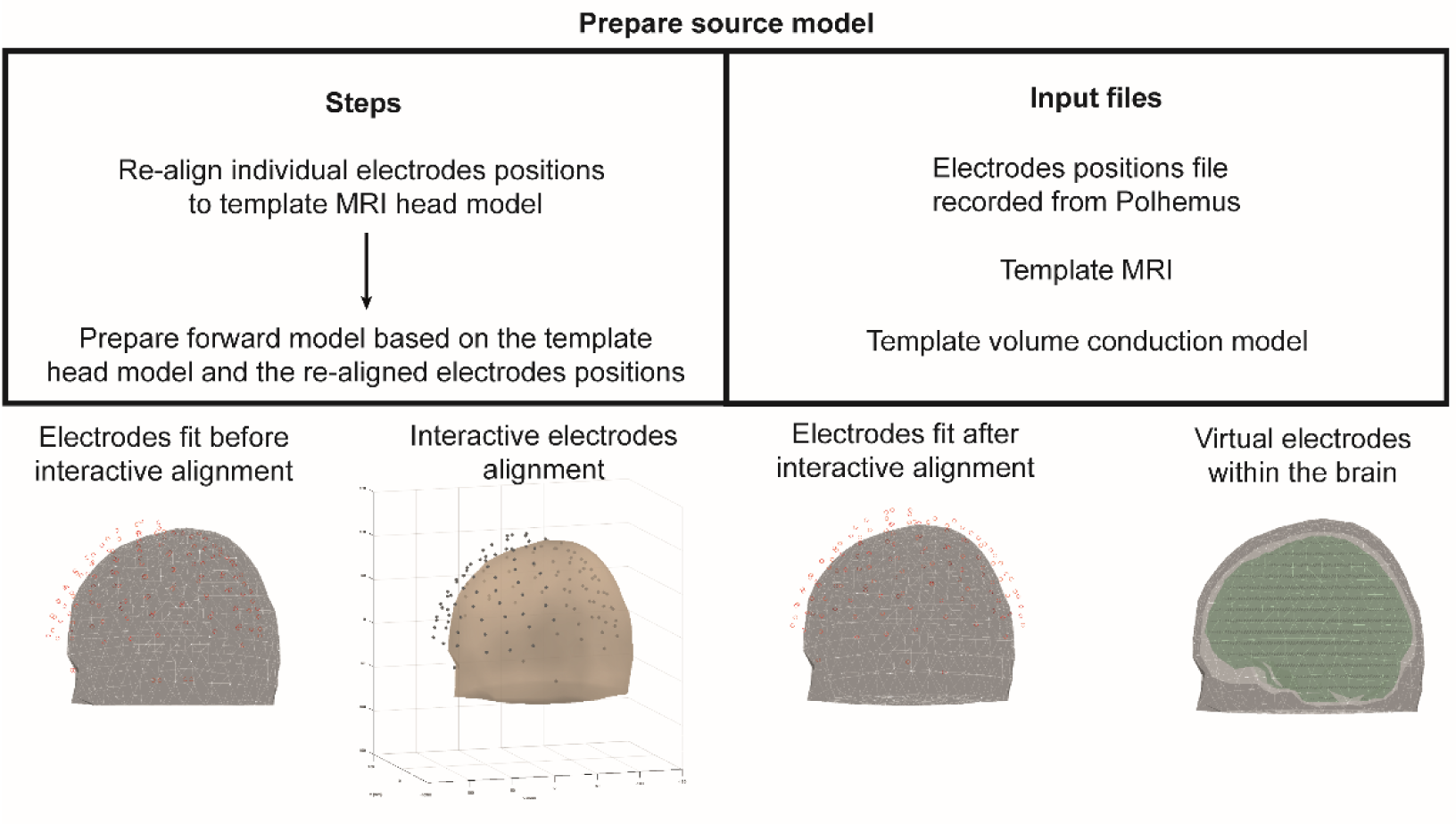
Source reconstruction pipeline part 1 – preparing the source model.

**Fig. S2:**
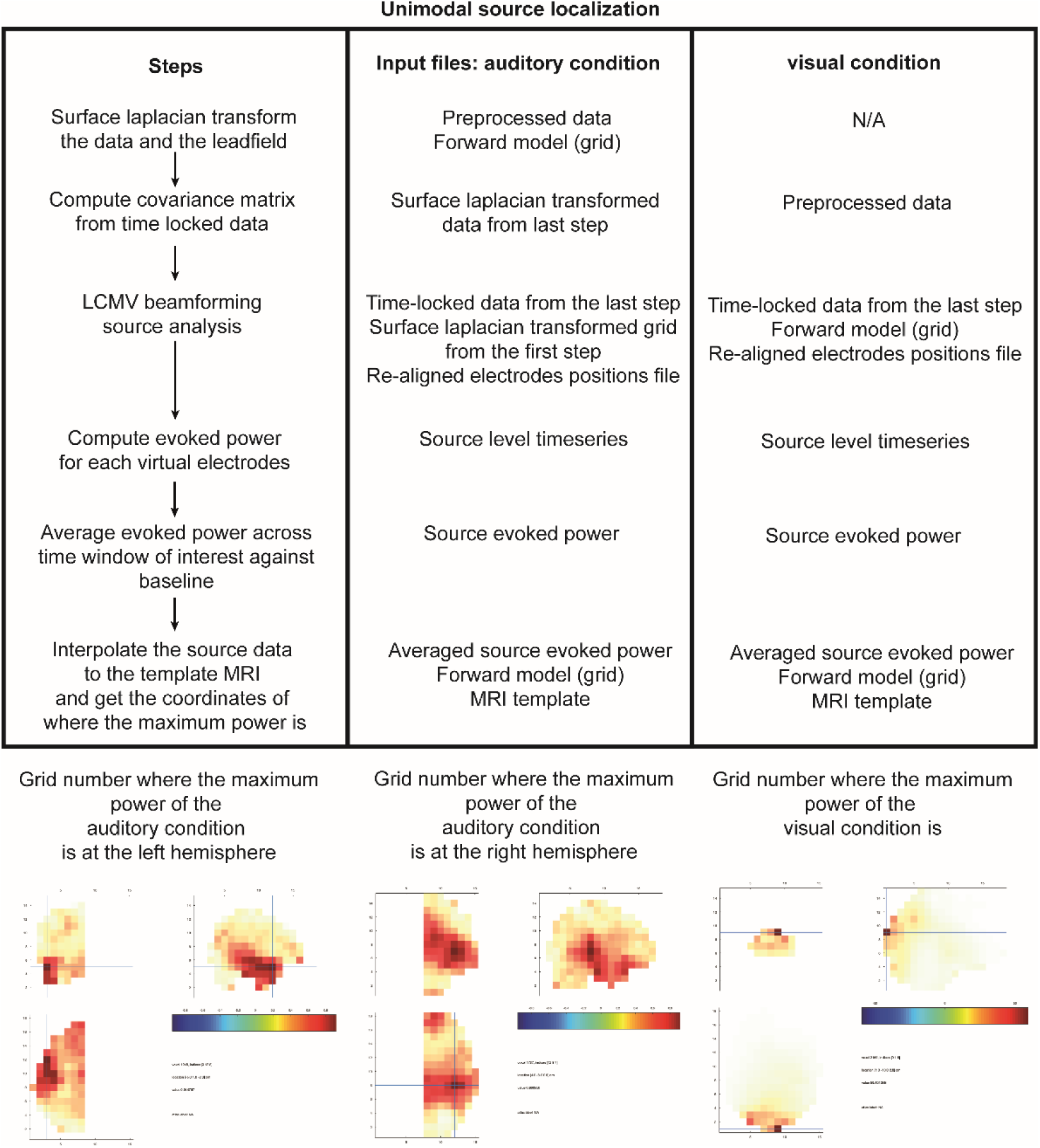
Source reconstruction pipeline part 2 – unimodal source localization.

**Fig. S3:**
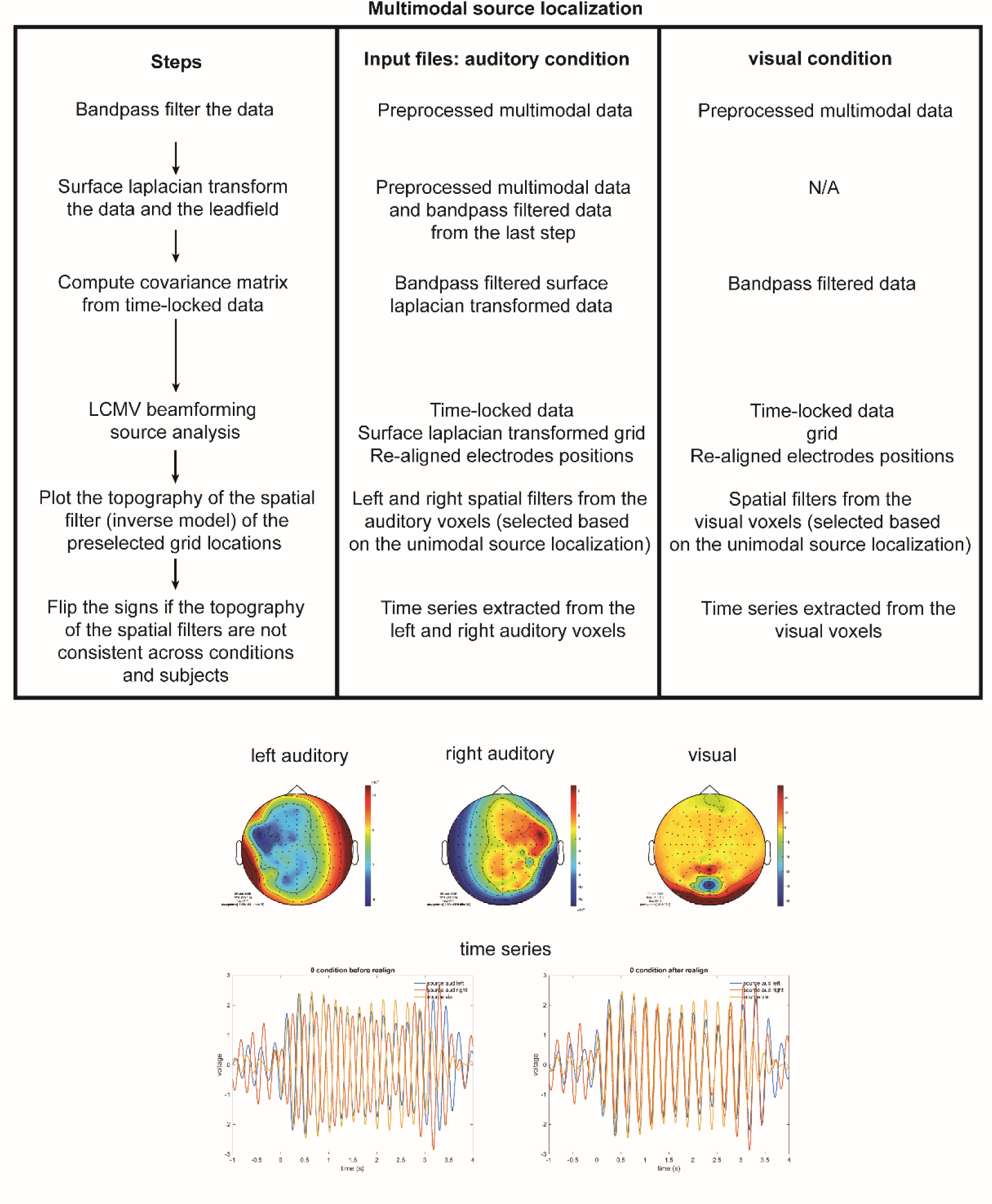
Source reconstruction pipeline part 3 – multimodal source reconstruction.

